# Modulation of locomotion and motor neuron response by the cohesive effect of acute and chronic feeding states and acute d-amphetamine treatment in zebrafish larvae

**DOI:** 10.1101/2024.01.28.577634

**Authors:** Pushkar Bansal, Mitchell F. Roitman, Erica E. Jung

## Abstract

Amphetamine (AMPH) increases locomotor activities in animals, and the locomotor response to AMPH is further modulated by caloric deficits such as food deprivation and restriction. The increment in locomotor activity regulated by AMPH-caloric deficit concomitance can be further modulated by varying feeding schedules (e.g. acute and chronic food deprivation and acute feeding after chronic food deprivation). However, the effects of different feeding schedules on AMPH-induced locomotor activity are yet to be explicated. Here, we have explored the stimulatory responses of acutely administered d-amphetamine in locomotion under systematically varying feeding states (fed/sated and food deprivation) and schedules (chronic and acute) in zebrafish larvae. We used wild-type and transgenic[Tg(mnx1:GCaMP5)] zebrafish larvae and measured swimming activity and spinal motor neuron activity *in vivo* in real-time in time-elapsed and cumulative manner pre- and post-AMPH treatment. Our results showed that locomotion and motor neuron activity increased in both chronic and acute food deprivation post-AMPH treatment cumulatively. A steady increase in locomotion was observed in acute food-deprivation compared to an immediate abrupt increase in chronic food-deprivation state. The ad libitum-fed larvae exhibited a moderate increase both in locomotion and motor neuron activity. Conversely to all other caloric states, food-sated (acute feeding after chronic food deprivation) larvae moved moderately less and exhibited a mild decrease in motor neuron activity after AMPH treatment. These results point to the importance of the feeding schedule in modulating amphetamine’s characteristic stimulatory response on behavior and motor neurons.

## INTRODUCTION

Amphetamine (AMPH) is a known psychostimulant drug and its chronic use results in its compulsive abuse which is one of the primary causes of AMPH addiction and its tolerance(Hyman, 1996; BERMAN et al., 2009). Rewarding and reinforcing effects of amphetamine play a major role in increasing the abuse liability of the drug and lead to an increment in physiological output in humans and animals, such as increased locomotor activity. A clinical study showed an increased locomotor output in humans post-amphetamine administration(Minassian et al., 2016). In preclinical studies using animals, long-term AMPH administration caused a progressive increase in motor activity in rats(Segal and Mandell, 1974).

Also, acute AMPH treatment in mice and pigeons resulted in increased locomotor activity and stereotypical behavior(Idemudia and McMillan, 1984; Yates et al., 2007). These changes in motor responses arose from low to moderate doses of amphetamine whereas AMPH intoxication has led to severe physiological responses behavior in animals and humans(Shiorring, 1980; Fitzgerald and Bronstein, 2013).

The motor response to amphetamine can be affected by the physiological state of the animal, such as hunger and satiety. Specifically, food deprivation/restriction is known to enhance the motor response to amphetamine. In preclinical studies, Monkeys exhibited an increased amphetamine intake in the food-restricted state which potentiated amphetamine’s stimulating effects by increasing their motor activity (Carroll et al., 1979). Similarly, food-restricted rodents showed enhanced rewarding effects of amphetamine by increasing their locomotion(De Vaca and Carr, 1998). Other studies involving food-restricted rodents showed an increased amphetamine intake along with higher conditioned place preference(Stuber et al., 2002; Geuzaine et al., 2014). Acute food restriction for 24 hours in rats altered locomotor activity when treated with amphetamine(Mabry and Campbell, 1975; Honma et al., 1992). In these pre-clinical studies, amphetamine-food restriction mediated interactive response was observed when animals were subjected mostly to chronic food-restriction state and few in acute food-restricted state in separate independent studies. To date, no study has systematically varied the caloric state of the animal by varying the feeding schedule while subjecting them to a caloric deficit state, such as acute/chronic food deprivation and satiation acute/ad libitum feeding, to investigate the behavioral and neural motoric effects of amphetamine.

Zebrafish larvae provide a good model organism to study motor behavioral and neural responses to addictive drugs. Zebrafish exhibit robust locomotor behavior and importantly, their optical transparency allows researchers to record and quantify individual motor neuron activity in the spinal cord using genetically encoded neural activity indicators(Muto et al., 2011). Larval zebrafish have shown behavioral sensitivity similar to mammals to a variety of addictive drugs including amphetamine(Irons et al., 2010; Cousin et al., 2014; Basnet et al., 2019). A recent study showed the effect of chronic food restriction on physiological responses, locomotion, heart rate, and motor neuron response(Bansal et al., 2023). Here, we address whether acute feeding and acute deprivation can differently modulate the motor response to amphetamine especially relative to the ad libitum fed and chronically deprived state. In zebrafish larvae, we measured the change in response in behavior and motor neurons in the spinal cord, an underlying neural substrate responsible for motor action, affected by amphetamine-caloric stress interaction. To quantify this, we analyzed locomotor and motor neuron activity in time elapsed manner and cumulatively for four different feeding conditions – Ad Libitum (unrestricted food availability), Food Sated (fed 24 hours before experimentation), Chronic Food Deprivation (never fed) and Acute Food Deprivation (24-hour starvation). Our findings showed that chronic and acute food deprivation potentiated the motor-stimulating effects of amphetamines in zebrafish. Both acute and chronic food deprivation induced a longer-lasting AMPH-mediated motor effect where acute food deprivation exhibited progressive increment in activity over intervals, unlike chronic food deprivation. Whereas motor response to AMPH in ad libitum and food-sated states was not significant. Overall, our study uncovers how systemic variation in caloric states evokes motor responses by distinctly modulating amphetamine’s inherent stimulatory response.

## Materials And Methods

Adult wild-type (abwt) and transgenic tg(mnx1:GCaMP6) zebrafish were housed at the zebrafish facility. They were maintained in an automated water racking system (Aquaneering, Inc. San Diego, CA). Approval for all conducted experiments was acquired from AALAC and all guidelines were diligently followed. The Zebrafish facility was maintained with a 14/10-hour light cycle schedule and was set at 28 deg C. For all experiments, 7 days post fertilization (dpf) zebrafish larvae were used. This study involves different feeding schedules to measure larval responses and zebrafish larvae develop neurons responsible for mediating feeding responses(Wee et al., 2019). Also, in the larval stage, this teleost has been shown to stay healthy without any phenotypical anomaly till 8dpf(Hernandez et al., 2018). Moreover, the feeding schedule organized for food-sated and acute food deprivation only allowed us to use 7dpf larvae for uniformity among all feeding states, thus, 7dpf seemed to be appropriate to conduct experiments. Further, the zebrafish rack system was also maintained at 28 degrees c temperature with water-conductivity between 600-750µs and pH was maintained within a range of 7.2-8.0. Larvae in the embryonic stage were initially kept in water with aforementioned conditions in the fish room in all states (harvesting, hatching, experiments) and were brought to the research facility when they turned 7 dpf.

### Experimental groups

We divided zebrafish into two groups namely control and AMPH_treatment groups which was further subdivided into eight groups, depending on their feeding states[Control: AL-AMPH (Ad libitum fed without AMPH), FS+AMPH (Food-sated with AMPH), CFD-AMPH (Chronic food-deprivation without AMPH), AFD-AMPH (Acute food-deprivation without AMPH), FS-AMPH (Food-sated without AMPH) and Treatment: AL+AMPH (Ad libitum fed with AMPH), CFD+AMPH (Chronic food-deprivation with AMPH), AFD+AMPH (Acute food-deprivation with AMPH)]. In the AL state, food was available ad libitum when they turned 5 dpf and remained available until the experiments were started. FS larvae were deprived of food until they turned 6 dpf and food was made available to them until they turned 7 dpf till the start of the experiment. Food was available to them for 24 hours. The larvae in the CFD state were deprived of food since they were born and were never fed at any point before the experiment. AFD larvae were fed till 6 dpf and larvae were transferred to a dish without food 24 hours before the experiments. A 24-hour food deprivation is called Acute food deprivation (Fig. 1A). Recording of locomotor activity and motor neuron activity was performed in both groups. These groups were further treated with a dose of 0.7uM of amphetamine (detailed procedure mentioned below). Previous studies performed in zebrafish larvae involving treatment with AMPH demonstrated a dose-dependent behavioral response that showed a ‘U’ pattern where locomotor activity decreased in low and high doses and increased in the moderate dose (0.7uM)(Irons et al., 2010). Hence, the finding of this and our previous study was the reason to select 0.7uM as a dose of choice where it would be interesting to see how different caloric states alter this significant motor response.

**Fig. 1.**
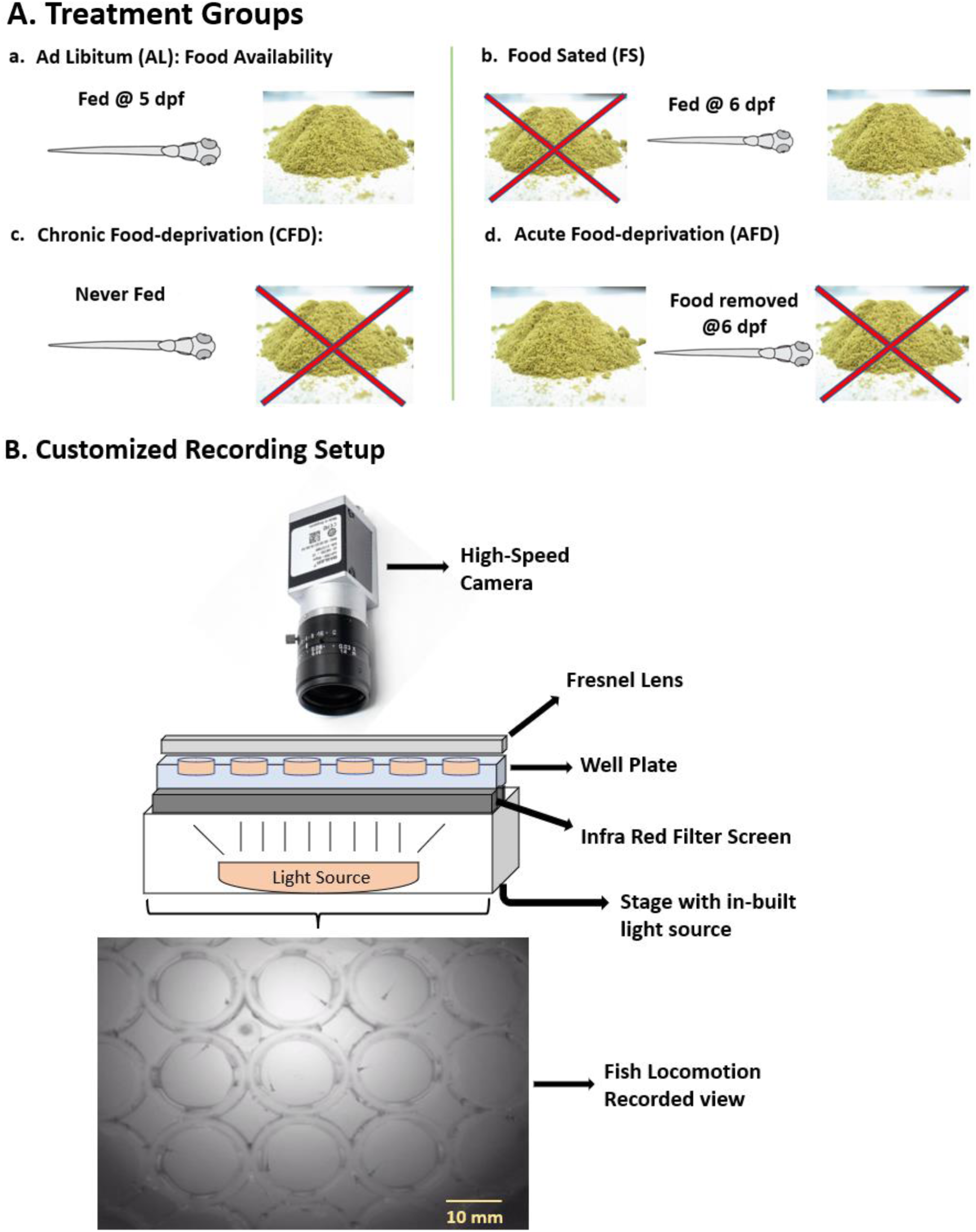
Experimental setup and treatment groups. **A.)** We used four different caloric states namely, Ad libitum (AL), Chronic food deprivation (CFD), Acute food deprivation (AFD) and Food-sated (FS). All larvae used here were 7 dpf at the time of the experiment. **a.)** In the Ad libitum fed (AL) state, food was available to AL larvae when they turned 5 dpf and remained available until the experiments commenced. **b.)** In the FS state, larvae were deprived of food initially and food was only made available to them when larvae turned 6dpf remained available till the start of the experiment. Food was available to them only 24 hours before the experiment is termed as food-sated state. **c.)** In the CFD state, larvae were deprived of food since they were born and were never fed at any point before or during the experiment. **d.)** In AFD state feeding started at 5 dpf where food was removed when larvae turned 6 dpf and were transferred to a dish without food 24 hours before the experiments. Here, 24-hour food deprivation is termed as Acute food deprivation. **B.)** This customized setup consisted of an invertedly placed high-speed camera that records the locomotor activity of zebrafish larvae placed in the well plate. An IR filter screen was placed between the plate and the light source to only IR light to pass and block all other wavelengths. The well plate was placed on the top of the in-built light source of a dissection microscope with an eyepiece and upper light source removed for easy recording. The plate was covered with a Fresnel lens to minimize field distortion.

### Locomotor activity setup and post-processing

For locomotion measurement and analysis, wild-type (abwt) larvae were used. In Fig. 1B, to construct a customized setup, we used a dismantled part of the dissection microscope that consisted of an in-built light source. An Infrared (IR) filter screen (43954, Edmund Optics, NJ, USA) was placed on the light source to allow only IR frequency light to pass through. Zebrafish larvae exhibited hyper-locomotion in the dark environment which was measured using IR light and larvae did not adversely affect fish behavior(Basnet et al., 2019). Thus, using IR light to mimic the dark environment would be interesting to examine its increased response in all four states and with amphetamine treatment. On the IR filter, we placed a 48-well plate in which larvae were placed individually in each well. To minimize distortion during recording activities from the wells from the corners, a 6.7” x 6.7” Fresnel lens (46614, Edmund Optics, NJ, USA) was placed over the 48-well plate. The whole setup was recorded by an invertedly placed high-speed camera (Basler Gencam1 GigE camera, 4.5 – 12mm, IR ½”). Activity recording was analyzed and generated using Ethovision XT16 (Noldus Information Technology, VA, USA). Generated data was binned within the software followed by statistical analysis.

Initially, both -AMPH and +AMPH groups were recorded for two separate 10-minute epochs (Baseline, Treatment). Before recording, larvae were allowed to acclimatize to the well environment for 5 mins. The baseline epoch in -AMPH groups were recorded for 10 mins in fresh fish water. After this recording period, the fish water was pipetted out and replaced by fish water and activity was recorded for another 10 mins (treatment epoch for -AMPH group). Similar to the -AMPH group, the baseline epoch was in a similar time and manner for 10 mins. It was followed by pipetting out the fish water and pipetting in 0.5ml of 0.7uM solution of d-amphetamine hemisulfate (Sigma Aldrich, MO). A period of 2-3 mins was permitted to larvae for the drug consumption in the well followed by another 10 min recording period (treatment epoch for +AMPH group). Further, each epoch is binned into five 2-min intervals and comparisons between binned epochs were made.

### Motor neuron activity measurement and post-processing

Motor neuron activity was measured in a transgenic zebrafish line tg(mnx1:GCaMP5) that expressed fluorescence in spinal motor neurons. Larvae were immobilized using 30ul of 300uM Pancuronium bromide solution (P1918, 10MG, Millipore Sigma, WI, USA), a paralyzing agent to inhibit any voluntary or involuntary activity and contraction for stable neuron recording. This procedure was followed by complete immobilization of the larvae in a 1.5% agarose gel drop. Gel drop is covered with fish water/amphetamine according to epochs (Baseline, Treatment) in -AMPH and +AMPH groups and recording manner and periods remained the same as in locomotor activity.

To record motor neuron activity, a wide-field inverted epifluorescence microscope was used. The microscope setup was connected to an X-cite 120 fluorescence illumination/excitation lamp setup (Excelitas Technologies Corp., Canada). GCaMP5 is a genetically encoded calcium indicator (GECI), and it has an illumination/excitation wavelength of 488nm/510nm, thus, a GFP bandpass filter (Chroma Technology Corp, VT) was installed to obtain the required excitation wavelength. A 40x water immersion objective (Olympus Corporation, Japan) was used through which the excitation light passes and was also used to image the fluorescence-emitting motor neurons. To record the spiking this neuron spiking activity, a Hamamatsu Orca Flash 4.0 high-speed sCMOS camera (Hamamatsu Photonics, Japan) at 35 fps. The whole recording setup was connected to HCImage Live, a recording and visualization software (Hamamatsu Photonics, Japan) and was used to record and export the recordings in a tif file format. Tif file frames were opened using ImageJ visualization and analysis software (National Institute of Health). Regions of Interest (ROIs) were selected using a polygon selection tool around the soma region of the neurons and their respective frame vs intensity data were generated within ImageJ. The data were exported to Microsoft Excel where frames were converted to time and this time vs intensity data was further exported into Origin Pro (Version 2023, OriginLab Corporation, Northampton, MA, USA). Data were smoothened using an FFT filter with window points=30. From the smoothed data, the baseline was calculated using an in-built function called ‘Peak Analyzer’ followed by the ‘subtract baseline’ option with customizable parameters. The obtained baseline was subtracted from smoothed intensity data and divided by baseline to obtain normalized data. Using the peak analyzed option, the find peaks option was used along with a height threshold of 5% to get peak information. The ‘peak info’ option was used to get the peak data (x-axis: time points, y: normalized intensity). These peak data were exported for statistical analysis.

### Statistical Analysis

Locomotor data were analyzed using GraphPad Prism (version 9.5.1, San Diego, CA, USA). Binned data were analyzed using one-way ANOVA with repeated measures followed by Dunnet’s multiple comparisons post-hoc tests. Cumulative data were analyzed using Two-way and Three-way ANOVA with repeated measures and Sidak and Tukey’s post-hoc tests were used respectively for multiple comparisons. For motor neuron analysis, inter-spike interval and ROC curves were analyzed using R programming and Python. Raster plots were created using Origin Pro (Version 2023, OriginLab Corporation, Northampton, MA, USA). Statistics were performed with * p<0.05, **p<0.01, ***p<0.001, p<0.0001 and non-significant relationships were not shown with any indication. Bars were represented with mean±SEM.

## RESULTS

### Food deprivation and satiation conversely alter locomotor activity post-AMPH treatment

We analyzed the time-elapsed swimming (locomotion) activity in all four caloric states and compared a 2 min control interval (fish water only) with amphetamine-treated intervals (five intervals each of 2mins) (Fig. 1A). Locomotion analysis was performed using RM One-Way ANOVA with intervals as a repeated measure. In the AL state, Two-way ANOVA did not reveal a significant interval effect (F(5,55) = 1.589, p>0.05) between intervals (Fig. 2A, a). The post-hoc comparison also did not indicate any significance between intervals post-AMPH administration. Similarly, in the FS state, the statistical test did not show a significant effect of interval on swimming activity after AMPH treatment (F(5,60) = 1.589, p>0.05).

**Figure 2.**
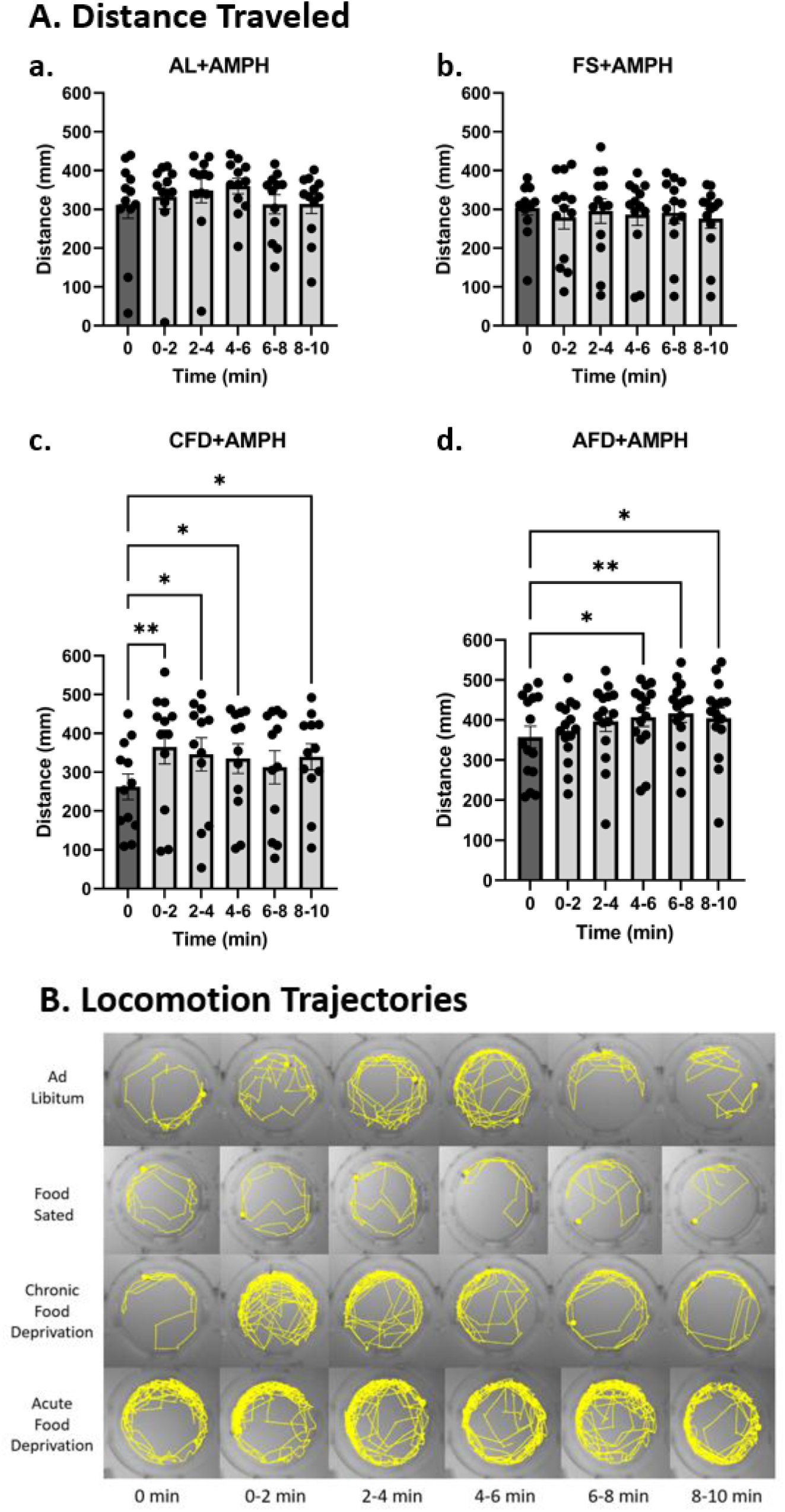
Time-elapsed locomotion response to amphetamine. A.) The time elapsed effect of amphetamine was assessed in binned 2 min intervals (10 mins in total) and all intervals were compared to a two min interval recorded before the addition of amphetamine. The initial 0 min interval is a 2 min long control interval without amphetamine (dark bar) and the rest (light bars) are amphetamine treated 2 min intervals which are compared to the control interval. a.) Distance increase observed in the AL state after AMPH was mild but non-significant in all five intervals compared to the control (0 min) interval (p>0.05). b.) FS state showed a mild non-significant decrease in distance traveled by larvae after AMPH treatment compared to a 2-min control interval (p>0.05). c.) Larvae in the CFD state showed the most significant increase instantaneously after AMPH treatment in 0-2 intervals (p<0.01) and the locomotion remained significantly high in all intervals compared to the control interval (p<0.05). d.) AFD larvae showed a significant increase in locomotion after a 4-min delay (first 2 mins interval) and kept increasing till the end (p<0.01). B.) Visual representation of time-elapsed locomotion trajectories from fish swimming shows the path covered by larvae in the wells before and after drug treatment

Planned comparison results also remained non-significant between intervals (Fig. 2A, b). Contrarily, chronic food deprivation significantly increased swimming activity in subjects after AMPH treatment (F(5,55) = 3.279, p<0.05) and planned comparisons between intervals also revealed a statistical significance across all intervals where AMPH showed an immediate effect after treatment (Fig. 2A, c). Larvae in AFD state post-AMPH treatment showed the most significant effect of intervals among all states (F(5,70) = 3.734), p<0.01). Post-hoc comparisons indicated significance between intervals compared to the control interval and the drug induced a significant locomotor activity after a 4 min delay (Fig. 2A, d).

Visual representation in Fig. 2B also shows the swimming activity trajectories where an overall higher swimming activity in control interval can be observed in acute food deprivation followed by food-sated, ad libitum state and least locomotor activity can be seen in chronic food deprivation state. Whereas amphetamine treatment abruptly increases locomotion in chronic food deprivation and follows a relatively gradual decrease in the activity. Acute food deprivation showed a progressive significant increase in locomotion after amphetamine administration. Trajectories in AL larvae showed a short-term increase in activity which decreased at the last interval. Locomotion trajectory in the FS state showed an overall decreased locomotor activity relative to the control interval as well as all other states.

Cumulative locomotor activity was also assessed in -/+AMPH groups. In this analysis, we added absolute distance traveled values continually (adding values continuously beginning from the first interval (0-2 min interval) to the following interval till the end of epochs (Fig. 3A). The perpetual increment of locomotion shows an overall gradual decrease in AL state in -/+AMPH between epochs (Fig. 3A, a).

**Figure 3.**
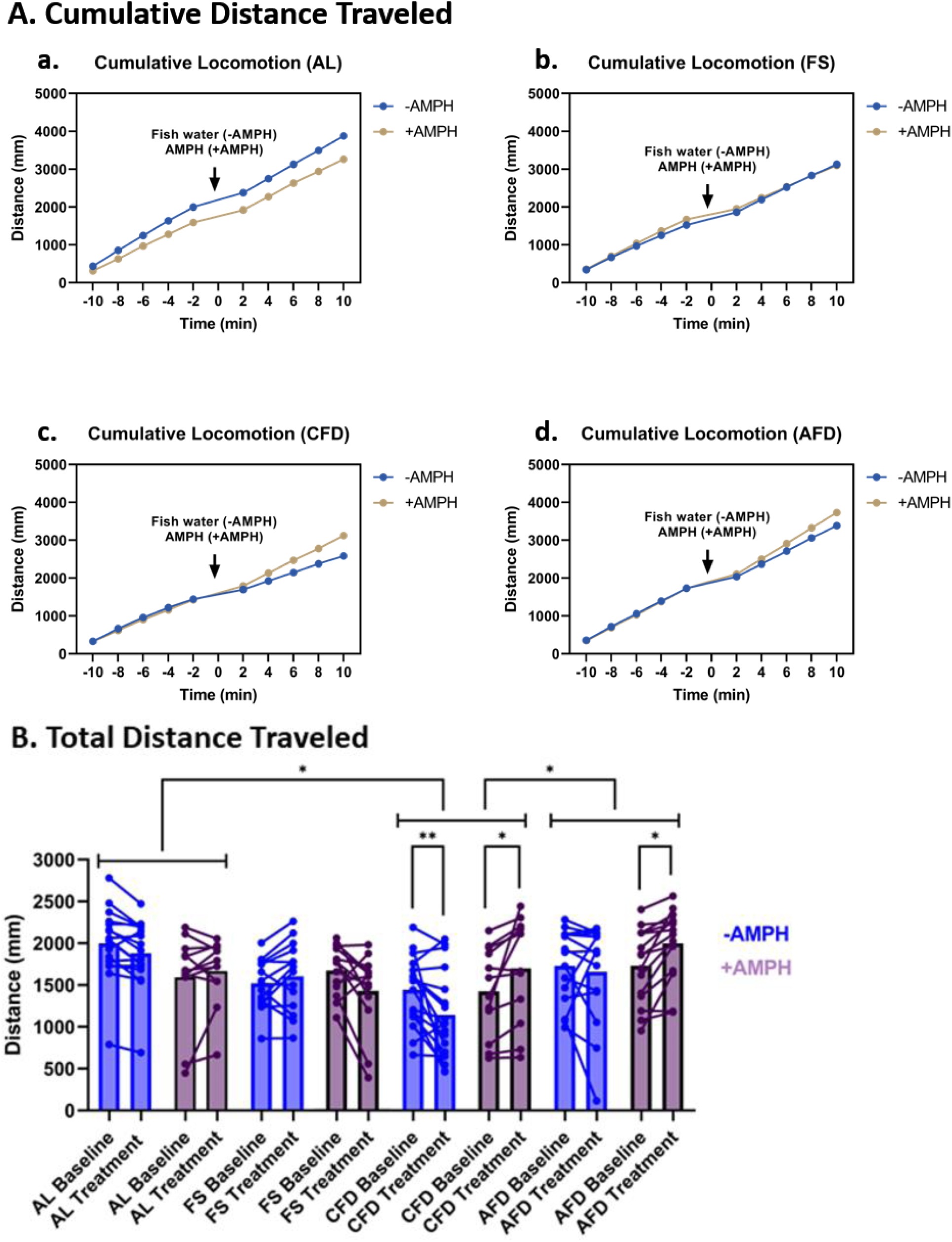
Time-elapsed cumulative and total distance traveled. A.) Distance traveled by fish in all states was measured cumulatively where distance traveled values in a previous interval are added to the subsequent interval values. a.) Cumulative locomotion of AL larvae in the -AMPH group traveled a longer distance than +AMPH in control and treatment epochs. b.) Larvae in the FS-AMPH group moved mildly longer distances cumulatively than FS+AMPH and AMPH treatment reverted the locomotion of both groups to an equal level. C.) CFD-AMPH group moved a similar distance in the baseline epoch and a sharp increase in locomotion occurred at the end of the treatment epoch. d.) Distance traveled by AFD+AMPH increased gradually in the treatment epoch compared to the AFD-AMPH group and locomotion in the baseline epoch in both groups remained the same. B.) Cumulative locomotion was analyzed for two separate 10-min epochs in both control and AMPH-treated groups in all four states using Three-way RM ANOVA (Two-way RM ANOVA for motor neuron activity) and comparisons were made between epochs (state x epoch x drug). Locomotor activity significantly changed between epochs in CFD in the control (p<0.01) as well as the AMPH-treated group (p<0.05). Cumulatively, control larvae moved significantly shorter distances whereas AMPH treated evoked significant hyperactivity between baseline and treatment epochs. AFD larvae only displayed a significant increase after AMPH treatment only between epochs. AL and FS failed to show any significance in control and AMPH treatment conditions. However, control FS larvae moved mildly longer distances and AMPH treatment caused a moderate non-significant decrease indicating FS state evokes hypoactive response to AMPH treatment opposite to CFD and AFD states.

Cumulative activity in the FS state slightly increased in -AMPH, however, AMPH treatment decreased the activity and the cumulative effect in -AMPH and +AMPH groups was equal (Fig. 3A, b). Cumulative activity increased in both food-deprivation regimes. The chronic food deprivation state showed the most obvious relative increment in between -/+AMPH groups (Fig. 3A, c) than the AFD state where the increase was slow and gradual (Fig. 3A, d). We also measured cumulative activity over 10-min epochs (Baseline and treatment) for all four states using Three-way RM ANOVA with state, epoch and drug being independent factors and locomotor activity as a dependent factor (Fig. 3B). In locomotor activity quantification we observed significant effects of state (F(3,108) = 4.791; p<0.01), epoch x drug interaction (F(1,108) = 10.27; p<0.01) and state x drug x epoch interaction (F(3,108) = 9.546; p<0.0001).

Post-hoc comparisons between epochs showed a significant decrease in the -AMPH group of CFD state (p<0.01) and a significant increase in +AMPH groups of CFD and AFD states (p<0.05). After amphetamine treatment, the post-hoc test showed a relative increase in locomotor activity in the AFD+AMPH state (p = 0.0121) than CFD+AMPH state (p = 0.0541) between epochs. State-wise planned comparison revealed an overall significant difference between AL/CFD states and CFD/AFD states (p<0.05) where state-wise effect between CFD/AFD state was relatively greater (p = 0.115) than AL/CFD (p = 0.118).

### AMPH treatment increases motor neuron activity response and spiking frequency in food-deprivation states

Previously, we analyzed the effects of caloric states and AMPH in locomotion (swimming activity). Phenotypic locomotor behavior is directly affected by spinal motor neurons which are influenced by numerous neuron interconnections to and from both identified and yet to be identified. Moreover, physiological states such as food deprivation and stimulant drugs such as amphetamines are known to affect specific neuron populations in the brain(Johansson et al., 2008; Daberkow et al., 2013; Alhadeff et al., 2019). Thus, to assess the cohesive effects of AMPH and feeding states we investigated the spinal motor neuron activity by recording the neuron spiking in four different aforementioned feeding states followed by acute amphetamine treatment. We explored the frequency of motor neuron spiking in terms of inter-spike interval activity. In Fig. 4, raster plots representing neuron spikes depict the distribution of spikes in all four states over 600 seconds of recording time before and after amphetamine treatment.

**Figure 4.**
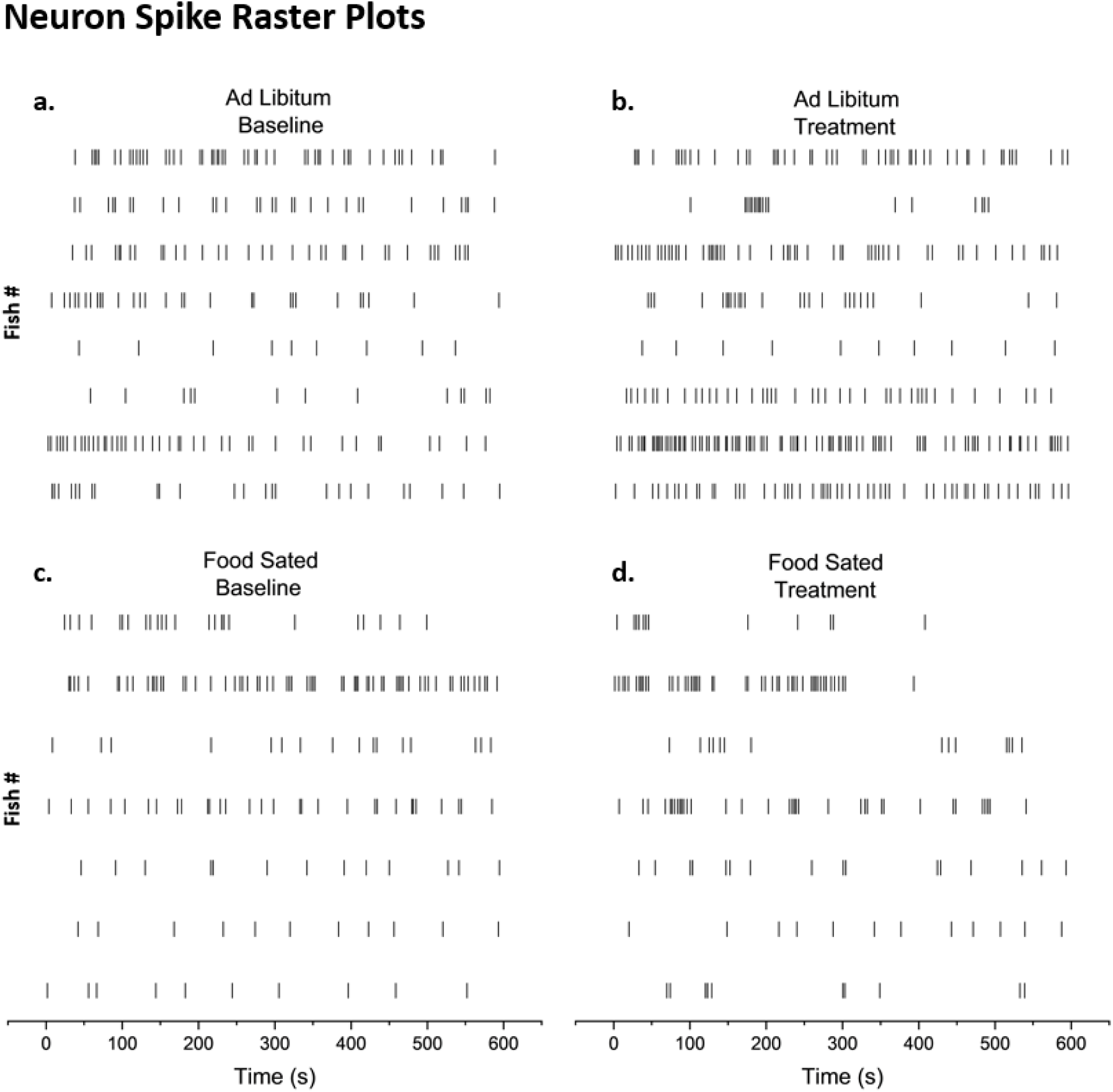
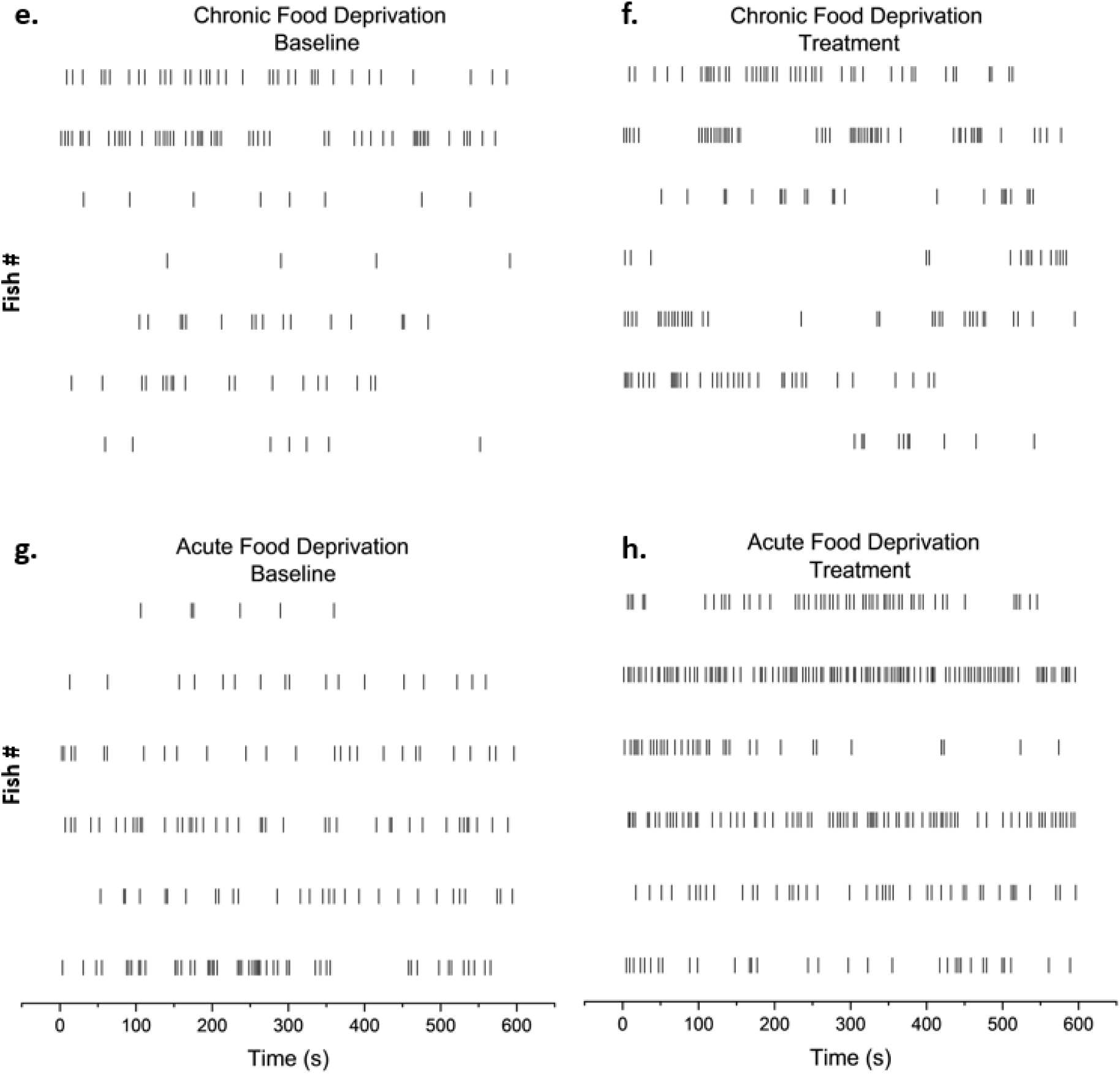
Raster plot representation for motor neuron distribution. Raster plots show the distribution of neurons spiking before and after amphetamine treatment. a-b.) Motor neuron activity in AL larvae mildly increased after amphetamine treatment. c-d.) Amphetamine although mildly decreased motor neuron activity in the FS state, overall change in activity remained non-significant. e-f.) Amphetamine induced a moderate increase in motor neuron activity. g-h.) Amphetamine evoked the most significant increase in motor neuron activity in the AFD state.

Information from these raster plots can also be used to extract useful information such as events with prolonged non-spike events, delays if any in amphetamine-induced effect, spike burst events etc. occurring in the neuron activity (Fig. 4A). To gain a deeper insight into inter-spike intervals (spike latency) pre- and post-AMPH treatment, we generated density plots between baseline(-AMPH) and Treatment (+AMPH) epochs in all four states (Fig. 5A).

**Figure 5.**
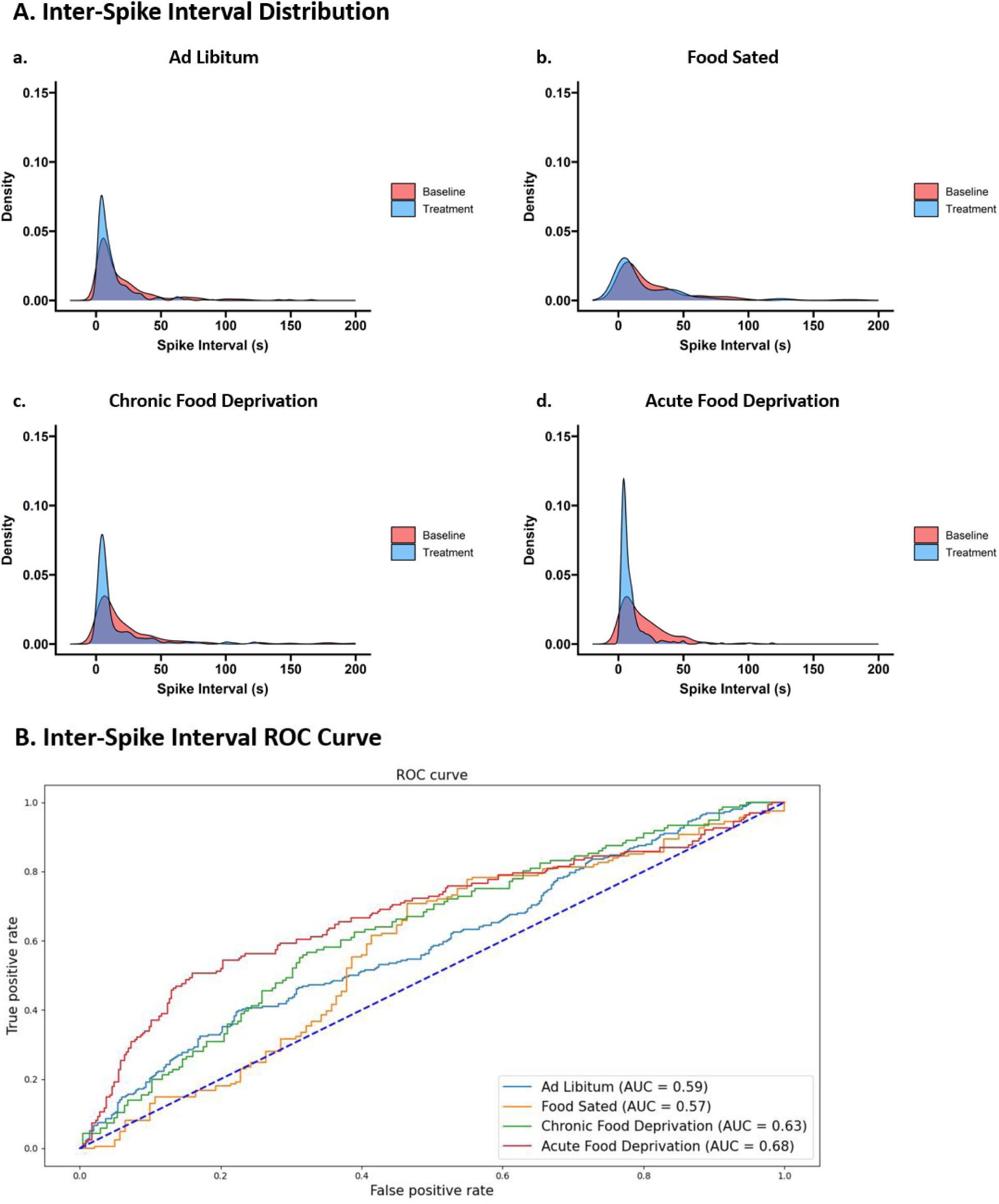
Inter spike interval analysis. Distribution and ROC curve of Latency (inter-spike interval) show the frequency of motor neuron spikes in all four caloric states and are compared between baseline and AMPH-treated groups. A.) Conditional density distribution of motor neuron inter-spiking frequency shows the effect of the drug on the latency in different feeding states. a.) Amphetamine treatment in AL larvae shows a mildly narrower distribution of spike latency compared to baseline. b.) Inter-spike interval distribution in FS state before and after AMPH treatment did not show a difference between the groups and has broadly spread distribution. c.) The latency distribution difference between baseline and treatment groups in the CFD state was moderate and distribution in the treated state was localized in smaller time intervals. d.) Inter-spike interval distribution was narrowest in the treated state compared to its baseline state as well as to all other states in all other groups. B.) The receiver operator characteristic (ROC) curve demonstrates the difference between neuron activity response in terms of inter-spike interval in all four states showing food-deprivation states potentiate the effect of amphetamine by increasing neuron spike occurrence rate (lowering inter-spike frequency) as compared to feeding states.

Density distribution in the Ad Libitum state shows a narrower interval distribution with a higher density peak at lower values after AMPH treatment with a mean interval value of 13.38 seconds and was mildly higher (19.06 seconds) before the treatment (Fig. 5A, a). Amphetamine treatment in food-sated larvae led to an average inter-spike interval time of 24.20 seconds which is higher than its control epoch i.e., 23.87 seconds indicating a minor decrease in occurrence of neuron activity post-AMPH administration.

Density peaks of control epochs are also higher than treatment epochs (Fig. 5A, b). Similar to AL larvae, CFD groups also showed an increase in density peak at lower interval values after AMPH treatment along with a decreased mean interval time (16.52 seconds) than baseline epoch (24.83 seconds) (Fig. 5A, c).

AMPH treatment in acute food deprivation (AFD) caused a remarkable change in the inter-spike interval by reducing it from 19.01 seconds in the baseline epoch to 10 seconds in the treatment epoch. The density peak at lower intervals was also noticeably higher after drug treatment (Fig. 5A, d). Overall, Amphetamine induced the highest change in inter-spike interval by an average of 9 seconds in AFD followed by the CFD group (8.3) seconds and AL group (5.8 seconds) whereas an opposite change was observed in the food-sated state (0.3 seconds In Fig. 5B, a detailed inter-spike interval analysis was performed using ROC curve method that compared the intervals between baseline and treatment epochs and calculating their specific areas under the curve (AUC) which represents the distinction in latency between baseline and treatment epoch in each state. In the AL state, AMPH treatment showed an area of 0.59 with p < 0.001 indicating a significant effect of the drug in decreasing latency of motor neuron spiking. A significant area under the curve of 0.57 was shown by the FS state (p<0.05) in response to the drug however, the drug in the FS state increased the latency. Chronic food-deprivation larvae exhibited an area of 0.63 with significance (p<0.0001) after baseline and treatment comparison. The highest and most significant area under the curve was shown by AFD larvae with an area of 0.68 (p<0.0001). ROC analysis distinguished the inter-spike interval between epochs significantly well in both food-deprivation regimes highly in the AFD state followed by the CFD state. Area measurement in both feeding states, AL and FS states was found to be significant however, the area under the curve was significantly lesser in the AL state followed FS state compared to food-deprivation states.

## DISCUSSION

Measurement of the integrated effect of caloric states-AMPH in terms of locomotor and motor neuron activity was performed here by acutely treating zebrafish larvae with d-amphetamine in four different caloric states. In this pre-clinical study, ad libitum (fully fed), caloric stress (chronic and acute food deprivation) and acute feeding (food-sated) states were investigated as potential alterants of stimulatory effects of d-amphetamine. While induction of psychostimulatory effects of AMPH has been documented in a handful of studies in food-restriction states (Deroche et al., 1993; De Vaca and Carr, 1998; Stuber et al., 2002; CABEZA DE VACA et al., 2004; Sharpe et al., 2012), behavioral and motor neuron activity outcomes of food reward (acute feeding) and amphetamine rewards in previous studies has only been observed when administered independently. Thus, the gap of knowledge in discerning the interactive outcome of acute feeding/food reward and amphetamine along with its comparison to other caloric states persists. Hence, this pre-clinical study investigates four different caloric states namely ad libitum. food-sated, chronic food-deprivation and acute food-deprivation states and strives to plug the knowledge gap by assessing pre-and post-AMPH mediated behavioral response (distance traveled) and motor neuron activity responses in aforenamed caloric states.

Our findings from comparing phenotypic locomotor behavior in all four states after AMPH treatment showed that both chronically and acutely food-deprived larvae displayed an increase in locomotion. In the Ad libitum state, larvae were slightly hyperactive (increased locomotor behavior) when AMPH was administered. Importantly, in a food-sated state, where larvae were acutely given access to food for 24 hours before the activity recording, decreased locomotor responses to AMPH treatment were observed. This novel observation expresses the acute food-rewarding effects on stimulatory effects of acutely administered amphetamine. These results support the observation in the studies cited above where sated animals were relatively less active than food-restricted animals after they were subjected to amphetamine treatment. Observations made in this study could be a consequence of a series of physiological and metabolic changes occurring inside the animal as a result of food stress and drug reward. For example, studies performed *in vivo* suggest that starvation causes muscles to utilize fatty acids (FA) as a primary energy source replacing ketones (Mehanna et al., 2008; Casper, 2020). Feeding FA supplements to rodents followed by amphetamine treatment resulted in significantly higher locomotor activity than in non-AMPH-treated counterparts (Trevizol et al., 2013). Food provided to zebrafish larvae in this study did contain oil (fish oil) which is a source of FA that was shortly removed in acute food-deprivation conditions and where metabolism of FA may have started only less than 24 hours ago. Additionally, fats tend to have a slower metabolism than other macronutrients resulting in retention of fats for longer periods. (Elia et al., 1999; Hargreaves and Spriet, 2020) Thus, although not immediate but gradual and long-lasting increase in locomotion observed in AFD larva could have been a result of FA metabolism commenced only a day before thereby leaving comparatively higher FA content in the fish than CFD fish and their interaction with d-amphetamine. The Yolk sac of zebrafish embryos has a significant amount of lipids and triglycerides which are converted into fatty acids (Sant and Timme-Laragy, 2018) and larvae break out of the yolk at 4dpf and remain without nutrition if not fed and CFD state may have led to a relatively higher FA metabolism due to extended starvation period depleting more FA in the CFD fish. Therefore, in CFD, where FA metabolism has been occurring for longer larval body might have caused an immediate increase in swimming activity after interacting with amphetamine.

For food-sated larvae, we hypothesized a spike in locomotor activity since both food and AMPH which are considered as rewards, both induce an increase in locomotion. Contrastingly, when food-sated larvae were exposed to amphetamine, their activity gradually decreased compared to the other three groups which could have been so due to the following reason. Zebrafish larva at the embryonic stage gets nutrients from the yolk and at 4dpf, the yolk gets dechorionated after which the larva depends on human feeding as mentioned before. Therefore, keeping the larva unfed and feeding it only 24 hours before the experimentation can be perceived as refeeding after a short period of starvation. In human subjects, after fasting, refeeding leads to a rapid decline in FA levels, lower than the fed baseline levels(Kolaczynski et al., 1996). A study performed in mice with an induced deficiency of omega-3 fatty acids followed by acute amphetamine treatment initially caused an increased locomotor activity but then a gradual decrease in locomotion was observed than control mice (McNamara et al., 2008).

Although fish food did have some fat content, metabolic changes arising from refeeding may have an overpowering effect in rapid FA declination. Larvae used in this study were already fed 24 hours before the activity recording. Thus, the aforementioned steep decline in FA levels lower than normal along with its interaction with amphetamine might be associated with the decrease in locomotor activity in acutely food-sated larvae. Future studies involving the fatty acid-AMPH interaction in different caloric states could shed light on the cause of this outcome by studying the behavioral responses and metabolism in zebrafish larvae and other animal models.

Agouti-related peptide (AgRP) neurons, a hunger-mediating neuronal population, get activated during starvation and cause increased locomotion activity (Krashes et al., 2011). A recent study shows that astrocyte expression is increased by the food-deprivation state and its coverage in AgRP neurons (Varela et al., 2021). Another research indicated that an increase in locomotor activity in amphetamine-treated mice requires the presence of astrocytes in the brain(Corkrum et al., 2019). Thus, concomitant expression of astrocytes and AgRP arising from food deprivation and additional exposure to AMPH could be a contributor to increased locomotor response in food-deprived larvae. In addition to AgRP neuron activation, acute starvation (AFD) increases food-seeking behavior (Ngo et al., 2022). In AFD larvae, a relatively prolonged increase in locomotor activity observed here post-amphetamine exposure could have been a result of both increased food-seeking and activity of AgRP neurons in acute removal of food than in CFD state, where AgRP activity could have faded due to chronic absence of food. Additionally, NPY neuron populations which are present in conjunction with AgRP neurons also get activated during CFD reducing energy expenditure and thereby reducing locomotion. (Luo et al., 2011; Paes and Fernandez, 2016) and NPY mRNA expression (required for NPY protein formation in NPY neurons) is reduced by amphetamine. (Anon, n.d.; Hsieh et al., 2005)) Although AgRP and NPY populations in zebrafish are expressed in distinct neurons, they are involved in appetite regulation. (Jeong et al., 2018) Hence, amphetamine-mediated reduction of NPY activity might have led to an uninhibited energy expenditure followed by increased locomotor activity. Locomotor activity in rodents with ablated POMC (hunger-suppressing neuron population) gene (Zhan et al., 2013) and amphetamine inhibits POMC neuron activity (Alhadeff et al., 2019). Both AgRP and POMC neuron populations are present and functional in zebrafish larvae (Zhang et al., 2012). Thus, in food-sated larvae, short-term activation of POMC neurons from acute feeding and sudden inhibition of POMC neurons from acute amphetamine treatment could be the reason for decreased swimming activity. Therefore, to fully understand the behavioral aspects of different caloric states, astrocyte-AgRP/NPY-POMC neuron nexus can be explored in future together with psychostimulants.

This study also analyses the change in motor neuron activity occurring in zebrafish larvae arising from amphetamine treatment in different feeding states. Our findings suggest that chronic and acute food deprivation states affected motor neuron activity the most after amphetamine treatment. We observed that the mean interval time between individual neuron spikes increased in food deprivation states than in fed states. Although the general functional connection between locomotion and motor neurons is known(Song et al., 2016; Gao et al., 2018), the findings presented here are novel in terms of recording the alteration in individual motor neuron signals in different feeding states as well as in response to the psychostimulant (AMPH) administration. Both acute and chronic food deprivation decreased POMC gene expression along with increased arcuate NPY mRNA expression observed only in chronic food deprivation and arcuate AgRP levels increased only in acute food deprivation. (Bi et al., 2003) Activated NPY neuron population relays inhibitory signals to the spinal cord (Duan, 2014; Koch, 2018). Study showing the effect of psychostimulants on NPY has not been performed extensively but one study shows that NPY levels decreased after amphetamine administration in rodents.(Kobeissy et al., 2008) Thus, expression studied with the drug may have caused NPY inhibition which significantly eliminated its inhibiting synapses towards the spinal cord resulting in its increase in motor neuron activity in chronically food-deprived zebrafish larvae. In the AFD state where AgRP neuron activity increases, occurs when ghrelin, which is a gut hormone released during the starvation state binds to AgRP receptors.(Cummings et al., 2001; Tschöp et al., 2001; Stuber et al., 2002; Bi et al., 2003; Müller et al., 2015) Study shows the presence of ghrelin receptors in the AgRP neuron population and ghrelin binding on these receptors increases the excitability of AgRP neurons(Chen et al., 2017). This AgRP activation sends synapses to Dopaminergic neurons via the Lateral hypothalamus in rodents(Castro et al., 2015; Khelifa et al., 2021). Furthermore, ghrelin receptors are also present on dopaminergic neurons in the Ventral Tegmental Area (VTA) in rodents(Quarta et al., 2009; Jerlhag et al., 2010; Edvardsson et al., 2021). Ghrelin injection in the VTA increased DA neuron activity in rodents which shows the direct action of ghrelin on DA neurons and DA populations send axonal projections to the spinal cord.(Abizaid, 2009; Reimer et al., 2013; Haehnel-taguchi et al., 2018) The Midbrain DA system responsible for rewarding effects in mammals is absent in zebrafish, however, research shows functional DA regions in zebrafish brain which have not been investigated in detail yet(Du et al., 2016; Barrios et al., 2020). Since amphetamine inhibits AgRP neuron activity and increases DA activity, future research could reveal if ghrelin binding on AgRP neurons is countering the inhibition of AgRP neurons and increasing the DA activity of known DA populations in zebrafish brain which is increasing the excitability of spinal motor neurons.

Food-sated (FS) larvae here showed a mild decrease in motor neuron activity post-AMPH treatment. FS state follows a pattern of starvation-feeding in this study. A recent study with similar fasting-refeeding showed an activation of a glutamatergic neuron population in the dorsomedial hypothalamus (DMH)(Imoto et al., 2021). Upon activation, these neurons project into the neurons of the periventricular nucleus of the hypothalamus (PVH) and PVH further projects into 5 different parts of the brain including the spinal cord (Ferguson et al., 2008; Geerling et al., 2010; Sutton et al., 2014). Future studies could explore these regions in response to acute food-sated state and amphetamine collectively to gain deeper insight into these responses. Involvement of arcuate nucleus has also been observed that is controlled by refeeding and feeding shortly after starvation abruptly increased POMC neuron activity and decreased NPY (Wu et al., 2014) and POMC neurons innervate. POMC neurons innervate the spinal cord and relay inhibitory signals.(Reinoß et al., 2020) Amphetamine decreases POMC neuron activity in rodents(Alhadeff et al., 2019) which should have increased spinal motor activity in normal ad libitum conditions. However, a sudden increase in POMC activity and an abrupt decrease in NPY along with acute amphetamine may have mildly decreased the motor neuron activity without showing a significant difference. Both POMC neuron populations and food-reward modulating dopaminergic populations are present in the hypothalamus in zebrafish brain and POMC possibly be projected into these hypothalamic DA neurons(Barrios et al., 2020; Reinoß et al., 2020) which can be studied in much detail in future. The effect of stimulants has not been studied to date in the aforementioned neuron populations in zebrafish. Altogether, these neuron populations along with the hormone ghrelin could be studied in future to obtain a complete understanding of the cohesive effect of caloric state and amphetamine on behavior and brain activity in this animal model. One of the main reasons for performing this study stems from the fact that feeding states and stimulants individually affect numerous neuron populations in the brain. Based on others and our results, the interconnections between neurons affected by caloric states and abusive drugs are fairly obvious(Anon, n.d.). Therefore, a thorough investigation of neural mechanisms underlying interactions between the neuronal groups will be beneficial for devising strategies to alleviate stimulant-mediated adverse effects.

## Conclusion

In conclusion, we investigated the change in stimulatory effects of acutely administered amphetamine in zebrafish larvae subjected to four different feeding states. We first measured the change in swimming behavior (distance traveled while swimming) in aforestated conditions. Our results show an increment in swimming distances traveled by both food-deprived larval groups which were abrupt but short-lived in chronic food-deprived state. Whereas a gradual prolonged increase in locomotor activity was observed in acute food-deprivation states after amphetamine was administered to them. Ad libitum and food-sated states had a modest effect on locomotion with a moderate activity increase in the former and a minor decrease in activity in the latter state post-AMPH treatment. Similarly, the spiking activity of spinal motor neurons was also analyzed in above stated conditions, and we observed a significant decrement in inter-spike intervals in acute food-deprivation and chronic food-deprivation states and a moderate increase in ad libitum state post-AMPH administration. AMPH treatment in food-sated larvae showed an unnoticeable increase in inter-spike latency. Overall, our findings show the significant potentiation effects of food-deficit states and a moderate attenuation effect of acute feeding (food-sated state) on amphetamine’s characteristic stimulatory effects on locomotor behavior and spinal motor neurons.

## Acknowledgements

We thank Dr. Claire Wyart for providing transgenic Tg(*mnx1*:GCaMP5) zebrafish line, Rachel for helping with data analysis and John for helping with maintenance of the fish facility.

## Author Contributions

**Erica E. Jung:** Conceptualization, Methodology, Writing-Reviewing and Editing, Funding acquisition, Project administration, Validation, Resources. **Pushkar Bansal**: Investigation, Data curation, Visualization, Formal analysis, Writing-Original draft preparation, Software. **Mitchell F. Roitman**: Writing-Reviewing and Editing, Data curation, Formal analysis, Validation, Software.

## Conflict of Interest

The authors have no conflict of interest to declare.

## Data Availability Statement

Data will be made available upon request to the corresponding author.

## Funding Sources

The work was supported by NARSAD Young Investigator Grant from the Brain & Behavior Research Foundation (BP and EEJ) and National Institute on Drug Abuse (Grant number: [DA050962: BP and EEJ] and [DA025634:MFR])

## Notes

### Competing Interest Statement

The authors have declared no competing interest.

### Summary of Updates

Updated the author order

## References

Abizaid A (2009) Ghrelin and Dopamine: New Insights on the Peripheral Regulation of Appetite. J Neuroendocrinol 21:787–793.

Alhadeff AL, Goldstein N, Park O, Klima ML, Vargas A, Betley JN (2019) Natural and Drug Rewards Engage Distinct Pathways that Converge on Coordinated Hypothalamic and Reward Circuits. Neuron 103:891–908.e6 Available at: 10.1016/j.neuron.2019.05.050.

Anon (n.d.) The Neuropeptides - Basic Neurochemistry - NCBI Bookshelf. Available at: https://www.ncbi.nlm.nih.gov/books/NBK28247/ [Accessed June 29, 2023a].

Anon (n.d.) A New Field of Neuroscience Aims to Map Connections in the Brain | Harvard Medical School. Available at: https://hms.harvard.edu/news/new-field-neuroscience-aims-map-connections-brain [Accessed June 23, 2023b].

Bansal P, Roitman MF, Jung EE (2023) Caloric state modulates locomotion, heart rate and motor neuron responses to acute administration of d-amphetamine in zebrafish larvae. Physiol Behav 264:114144 Available at: 10.1016/j.physbeh.2023.114144.

Barrios JP, Wang WC, England R, Reifenberg E, Douglass AD (2020) Hypothalamic Dopamine Neurons Control Sensorimotor Behavior by Modulating Brainstem Premotor Nuclei in Zebrafish. Curr Biol 30:4606–4618.e4 Available at: 10.1016/j.cub.2020.09.002.

Basnet RM, Zizioli D, Taweedet S, Finazzi D, Memo M (2019) Zebrafish larvae as a behavioral model in neuropharmacology. Biomedicines 7.

Berman SM, Kuczenski R, Mccracken JT, London ED (2009) Potential adverse effects of amphetamine treatment on brain and behavior: a review. Mol Psychiatry 14:123–142.

Bi S, Robinson BM, Moran TH (2003) Acute food deprivation and chronic food restriction differentially affect hypothalamic NPY mRNA expression. Am J Physiol - Regul Integr Comp Physiol 285:1030– 1036.

Cabeza De Vaca S, Krahne LL, Carr KD (2004) A progressive ratio schedule of self-stimulation testing in rats reveals profound augmentation of d-amphetamine reward by food restriction but no effect of a sensitizing regimen of d-amphetamine. Psychopharmacologia 175:106–113.

Carroll ME, France CP, Meisch RA (1979) Food Deprivation Increases Oral and Intravenous Drug Intake in Rats Published by : American Association for the Advancement of Science Stable URL : https://www.jstor.org/stable/1748263. 205:319–321.

Casper RC (2020) Might Starvation-Induced Adaptations in Muscle Mass, Muscle Morphology and Muscle Function Contribute to the Increased Urge for Movement and to Spontaneous Physical Activity in Anorexia Nervosa? Nutrients 12.

Castro DC, Cole SL, Berridge KC (2015) Lateral hypothalamus, nucleus accumbens, and ventral pallidum roles in eating and hunger: interactions between homeostatic and reward circuitry. Front Syst Neurosci 9:90.

Chen SR, Chen H, Zhou JJ, Pradhan G, Sun Y, Pan HL, Li DP (2017) Ghrelin receptors mediate ghrelin-induced excitation of agouti-related protein/neuropeptide Y but not pro-opiomelanocortin neurons. J Neurochem 142:512–520.

Corkrum M, Covelo A, Lines J, Thomas MJ, Kofuji P, Araque A, Corkrum M, Covelo A, Lines J, Bellocchio L, Pisansky M, Loke K, Quintana R, Rothwell PE, Lujan R, Marsicano G, Martin ED, Thomas MJ, Kofuji P (2019) Dopamine-Evoked Synaptic Regulation in the Nucleus Accumbens Requires Astrocyte Activity Article Dopamine-Evoked Synaptic Regulation in the Nucleus Accumbens Requires Astrocyte Activity. Neuron 105:1036–1047.e5 Available at: 10.1016/j.neuron.2019.12.026.

Cousin MA, Ebbert JO, Wiinamaki AR, Urban MD, Argue DP, Ekker SC, Klee EW (2014) Larval zebrafish model for FDA-approved drug repositioning for tobacco dependence treatment. PLoS One 9:e90467-.

Cummings DE, Purnell JQ, Frayo RS, Schmidova K, Wisse BE, Weigle DS (2001) A preprandial rise in plasma ghrelin levels suggests a role in meal initiation in humans. Diabetes 50:1714–1719.

Daberkow DP, Brown HD, Bunner KD, Kraniotis SA, Doellman MA, Ragozzino ME, Garris PA, Roitman MF (2013) Amphetamine paradoxically augments exocytotic dopamine release and phasic dopamine signals. J Neurosci 33:452–463.

De Vaca SC, Carr KD (1998) Food restriction enhances the central rewarding effect of abused drugs. J Neurosci 18:7502–7510.

Deroche V, Piazza PV, Casolini P, Le Moal M, Simon H (1993) Sensitization to the psychomotor effects of amphetamine and morphine induced by food restriction depends on corticosterone secretion. Brain Res 611:352–356.

Du Y, Guo Q, Shan M, Wu Y, Huang S, Zhao H, Hong H, Yang M, Yang X, Ren L, Peng J, Sun J, Zhou H, Li S, Su B (2016) Spatial and Temporal Distribution of Dopaminergic Neurons during Development in Zebrafish. Front Neuroanat 10:115.

Edvardsson CE, Vestlund J, Jerlhag E (2021) A ghrelin receptor antagonist reduces the ability of ghrelin, alcohol or amphetamine to induce a dopamine release in the ventral tegmental area and in nucleus accumbens shell in rats. Eur J Pharmacol 899:174039 Available at: 10.1016/j.ejphar.2021.174039.

Elia M, Stubbs RJ, Henry CJK (1999) Differences in fat, carbohydrate, and protein metabolism between lean and obese subjects undergoing total starvation. Obes Res 7:597–604.

Ferguson A V, Latchford KJ, Samson WK (2008) The paraventricular nucleus of the hypothalamus - a potential target for integrative treatment of autonomic dysfunction. Expert Opin Ther Targets 12:717–727.

Fitzgerald KT, Bronstein AC (2013) Adderall® (Amphetamine-Dextroamphetamine) Toxicity. Top Companion Anim Med 28:2–7 Available at: 10.1053/j.tcam.2013.03.002.

Gao S, Guan SA, Fouad AD, Meng J, Kawano T, Huang Y-C, Li Y, Alcaire S, Hung W, Lu Y, Qi YB, Jin Y, Alkema M, Fang-Yen C, Zhen M (2018) Excitatory motor neurons are local oscillators for backward locomotion. Elife 7.

Geerling JC, Shin J-W, Chimenti PC, Loewy AD (2010) Paraventricular hypothalamic nucleus: Axonal projections to the brainstem. J Comp Neurol 518:1460–1499.

Geuzaine A, Tyhon A, Grisar T, Brabant C, Lakaye B, Tirelli E (2014) Amphetamine reward in food restricted mice lacking the melanin-concentrating hormone receptor-1. Behav Brain Res 262:14–20 Available at: 10.1016/j.bbr.2013.12.052.

Haehnel-taguchi M, Fernandes AM, Böhler M, Schmitt I (2018) Projections of the Diencephalospinal Dopaminergic System to Peripheral Sense Organs in Larval Zebrafish (Danio rerio). 12:1–21.

Hargreaves M, Spriet LL (2020) Skeletal muscle energy metabolism during exercise. Nat Metab 2:817– 828 Available at: 10.1038/s42255-020-0251-4.

Hernandez RE, Galitan L, Cameron J, Goodwin N, Ramakrishnan L (2018) Delay of Initial Feeding of Zebrafish Larvae Until 8 Days Postfertilization Has No Impact on Survival or Growth Through the Juvenile Stage. Zebrafish 15:515–518.

Honma S, Kanematsu N, Honma KI (1992) Entrainment of methamphetamine-induced locomotor activity rhythm to feeding cycles in SCN-lesioned rats. Physiol Behav 52:843–850.

Hsieh Y-S, Yang S-F, Kuo D-Y (2005) Amphetamine, an appetite suppressant, decreases neuropeptide Y immunoreactivity in rat hypothalamic paraventriculum. Regul Pept 127:169–176.

Hyman SE (1996) Addiction to Cocaine and Amphetamine. Neuron 16:901–904.

Idemudia SO, McMillan DE (1984) Effects of d-amphetamine on spontaneous motor activity in pigeons. Psychopharmacology (Berl) 84:315–317.

Imoto D, Yamamoto I, Matsunaga H, Yonekura T, Lee M, Kato KX, Yamasaki T, Xu S, Ishimoto T, Yamagata S, Otsuguro K (2021) Refeeding activates neurons in the dorsomedial hypothalamus to inhibit food intake and promote positive valence. Mol Metab 54:101366 Available at: 10.1016/j.molmet.2021.101366.

Irons TD, MacPhail RC, Hunter DL, Padilla S (2010) Acute neuroactive drug exposures alter locomotor activity in larval zebrafish. Neurotoxicol Teratol 32:84–90 Available at: 10.1016/j.ntt.2009.04.066.

Jeong I, Kim E, Kim S, Kim HK, Lee DW, Seong JY, Park HC (2018) mRNA expression and metabolic regulation of npy and agrp1/2 in the zebrafish brain. Neurosci Lett 668:73–79 Available at: 10.1016/j.neulet.2018.01.017.

Jerlhag E, Egecioglu E, Dickson SL, Engel JA (2010) Ghrelin receptor antagonism attenuates cocaine- and amphetamine-induced locomotor stimulation, accumbal dopamine release, and conditioned place preference. Psychopharmacologia 211:415–422.

Johansson A, Fredriksson R, Winnergren S, Hulting A-L, Schiöth HB, Lindblom J (2008) The relative impact of chronic food restriction and acute food deprivation on plasma hormone levels and hypothalamic neuropeptide expression. Pept (New York, NY 1980) 29:1588–1595.

Khelifa MS, Skov LJ, Holst B (2021) Biased Ghrelin Receptor Signaling and the Dopaminergic System as Potential Targets for Metabolic and Psychological Symptoms of Anorexia Nervosa. Front Endocrinol 12:734547.

Kobeissy FH, Jeung JA, Warren MW, Geier JE, Gold MS (2008) Changes in leptin, ghrelin, growth hormone and neuropeptide-Y after an acute model of MDMA and methamphetamine exposure in rats. Addict Biol 13:15–25.

Kolaczynski JW, Considine R V, Ohannesian J, Marco C, Opentanova I, Nyce MR, Myint M, Caro JF (1996) Responses of leptin to short-term fasting and refeeding in humans: a link with ketogenesis but not ketones themselves. Diabetes 45:1511–1515.

Krashes MJ, Koda S, Ye CP, Rogan SC, Adams AC, Cusher DS, Maratos-Flier E, Roth BL, Lowell BB (2011) Rapid, reversible activation of AgRP neurons drives feeding behavior in mice. J Clin Invest 121:1424–1428.

Luo N, Marcelin G, Liu SM, Schwartz G, Chua S (2011) Neuropeptide Y and agouti-related peptide mediate complementary functions of hyperphagia and reduced energy expenditure in leptin receptor deficiency. Endocrinology 152:883–889.

Mabry PD, Campbell BA (1975) Potentiation of amphetamine-induced arousal by food deprivation: Effect of hypothalamic lesions. Physiol Behav 14:85–88.

McNamara RK, Sullivan J, Richtand NM, Jandacek R, Rider T, Tso P, Campbell N, Lipton J (2008) Omega-3 fatty acid deficiency augments amphetamine-induced behavioral sensitization in adult DBA/2J mice: Relationship with ventral striatum dopamine concentrations. Synapse 62:725–735.

Mehanna HM, Moledina J, Travis J (2008) Refeeding syndrome: what it is, and how to prevent and treat it. BMJ 336:1495–1498.

Minassian A, Young JW, Cope ZA, Henry BL, Geyer MA, Perry W (2016) Amphetamine increases activity but not exploration in humans and mice. Psychopharmacology (Berl) 233:225–233.

Müller TD, Nogueiras R, Andermann ML, Andrews ZB, Anker SD, Argente J, Batterham RL, Benoit SC, Bowers CY, Broglio F, Casanueva FF, Alessio DD, Depoortere I, Geliebter A (2015) Ghrelin. 4:437– 460.

Muto A, Ohkura M, Kotani T, Higashijima SI, Nakai J, Kawakami K (2011) Genetic visualization with an improved GCaMP calcium indicator reveals spatiotemporal activation of the spinal motor neurons in zebrafish. Proc Natl Acad Sci U S A 108:5425–5430.

Ngo F-Y, Li H, Zhang H, Lau C-YG (2022) Acute Fasting Modulates Food-Seeking Behavior and Neural Signaling in the Piriform Cortex. Nutrients 14:4156-.

Paes MR, Fernandez R (2016) Evaluation of energy expenditure in forward and backward movements performed by soccer referees. Brazilian J Med Biol Res 49:e5061–e5061.

Quarta D, Di Francesco C, Melotto S, Mangiarini L, Heidbreder C, Hedou G (2009) Rapid communication: Systemic administration of ghrelin increases extracellular dopamine in the shell but not the core subdivision of the nucleus accumbens. Neurochem Int 54:89–94.

Reimer MM, Norris A, Ohnmacht J, Patani R, Zhong Z, Dias TB, Kuscha V, Scott AL, Chen Y, Rozov S, Frazer SL, Wyatt C, Higashijima S, Patton EE, Panula P, Chandran S, Becker T, Becker CG (2013) Article Dopamine from the Brain Promotes Spinal Motor Neuron Generation during Development and Adult Regeneration. Dev Cell 25:478–491 Available at: 10.1016/j.devcel.2013.04.012.

Reinoß P, Ciglieri E, Minére M, Bremser S, Klein A, Löhr H, Fuller PM, Büschges A, Kloppenburg P, Fenselau H, Hammerschmidt M (2020) Hypothalamic Pomc Neurons Innervate the Spinal Cord and Modulate the Excitability of Premotor Circuits. Curr Biol 30:4579–4593.e7.

Sant KE, Timme-Laragy AR (2018) Zebrafish as a Model for Toxicological Perturbation of Yolk and Nutrition in the Early Embryo. Curr Environ Heal reports 5:125–133.

Segal DS, Mandell AJ (1974) Long-term administration of d-amphetamine: Progressive augmentation of motor activity and stereotypy. Pharmacol Biochem Behav 2:249–255.

Sharpe AL, Klaus JD, Beckstead MJ (2012) Meal schedule influences food restriction-induced locomotor sensitization to methamphetamine. Psychopharmacology (Berl) 219:795–803.

Shiorring E (1980) Psychopathology induced by “speed” drugs - (amphetamine, cocaine and related compounds). Aggress Behav 6:260 Available at: http://dx.doi.org/.

Song J, Ampatzis K, Björnfors ER, Manira A El (2016) Motor neurons control locomotor circuit function retrogradely via gap junctions. Nature 529:399–402 Available at: 10.1038/nature16497.

Stuber GD, Evans SB, Higgins MS, Pu Y, Figlewicz DP (2002) Food restriction modulates amphetamine-conditioned place preference and nucleus accumbens dopamine release in the rat. Synapse 46:83– 90.

Sutton AK, Pei H, Burnett KH, Myers Jr MG, Rhodes CJ, Olson DP (2014) Control of food intake and energy expenditure by Nos1 neurons of the paraventricular hypothalamus. J Neurosci 34:15306– 15318.

Trevizol F, Roversi K, Dias VT, Roversi K, Pase CS, Barcelos RCS, Benvegnu DM, Kuhn FT, Dolci GS, Ross DH, Veit JC, Piccolo J, Emanuelli T, Bürger ME (2013) Progress in Neuro-Psychopharmacology & Biological Psychiatry In fl uence of lifelong dietary fats on the brain fatty acids and amphetamine-induced behavioral responses in adult rat. Prog Neuropsychopharmacol Biol Psychiatry 45:215–222 Available at: 10.1016/j.pnpbp.2013.06.007.

Tschöp M, Wawarta R, Riepl RL, Friedrich S, Bidlingmaier M, Landgraf R, Folwaczny C (2001) Post-prandial decrease of circulating human ghrelin levels. J Endocrinol Invest 24:RC19–21.

Varela L, Stutz B, Song JE, Kim JG, Liu Z-W, Gao X-B, Horvath TL (2021) Hunger-promoting AgRP neurons trigger an astrocyte-mediated feed-forward autoactivation loop in mice. J Clin Invest 131.

Wee CL, Song EY, Johnson RE, Ailani D, Randlett O, Kim JY, Nikitchenko M, Bahl A, Yang CT, Ahrens MB, Kawakami K, Engert F, Kunes S (2019) A bidirectional network for appetite control in larval zebrafish. Elife 8.

Wu Q, Lemus MB, Stark R, Bayliss JA, Reichenbach A, Lockie SH, Andrews ZB (2014) The Temporal Pattern of cfos Activation in Hypothalamic, Cortical, and Brainstem Nuclei in Response to Fasting and Refeeding in Male Mice. Endocrinol 155:840–853.

Yates JW, Meij JTA, Sullivan JR, Richtand NM, Yu L (2007) Bimodal effect of amphetamine on motor behaviors in C57BL/6 mice. Neurosci Lett 427:66–70.

Zhan C, Zhou J, Feng Q, Zhang J-E, Lin S, Bao J, Wu P, Luo M (2013) Acute and long-term suppression of feeding behavior by POMC neurons in the brainstem and hypothalamus, respectively. J Neurosci 33:3624–3632.

Zhang C, Forlano PM, Cone RD (2012) Short Article AgRP and POMC Neurons Are Hypophysiotropic and Coordinately Regulate Multiple Endocrine Axes in a Larval Teleost. Cell Metab 15:256–264 Available at: 10.1016/j.cmet.2011.12.014.

